# Rapid task-dependent tuning of the mouse olfactory bulb

**DOI:** 10.1101/467530

**Authors:** Anzhelika Koldaeva, Andreas T Schaefer, Izumi Fukunaga

## Abstract

Adapting neural representation to rapidly changing behavioural demands is a key challenge for the nervous system. Here, we demonstrate that the output of the primary olfactory area, the mouse olfactory bulb, is already a target of dynamic and reproducible modulation. The modulation depends on the stimulus tuning of a given neuron, making olfactory responses more discriminable through selective amplification in a demand-specific way.

## Introduction

Behavioral contexts often pose conflicting demands on neural representations of stimuli. A representation that is optimal for one behavior may be unsuitable for another, as seen in the requirements for discriminating between stimuli vs. generalizing over the same stimuli. Adjusting representations transiently may enable the organism to cope with rapidly changing behavioral contexts.

Recent reports indicate that changes in sensory processing already in the primary olfactory region of the mouse, the olfactory bulb (OB), accompany behavioural acquisition of olfactory discrimination. For example, when observed over a period of several days, olfactory responses of OB output neurons change with learning (Chu et al., 2016; Yamada et al., 2017), though some of these changes appear to be associated with animals becoming familiar with the stimuli (Chu et al., 2016). Whether changes in the OB are dynamic, and how they are implemented remains unclear.

Here, we investigate how behavioural demands shape olfactory responses of OB output as mice switch between tasks that differ in difficulty. We find that olfactory processing in the OB in response to identical stimuli changes rapidly with task demands, in a manner that is suited to the task at hand.

## Results

To study how behavioural demands influence olfactory processing, we used two olfactory discrimination tasks, involving coarse versus fine discrimination (Fig. 1A). Olfactory stimuli used in the coarse discrimination task are easily distinguishable, while stimuli employed in the fine discrimination task are more similar (Supplementary Fig 1). In both tasks, the rewarded odor **α** (S+ odor) was a mixture of two odorants (A and B), mixed at a concentration ratio of 40%/60%. The nature of the task depended only on the non-rewarded odor (S-odor). In the coarse discrimination task, the S-odor was odor **β**, comprising odorants C and D, which are not present in the rewarded stimulus. On the other hand, the S-odor used in the fine discrimination task was odor **α^’^**, made by mixing odorants A and B, but mixed at a different concentration ratio (60%/40%). By using the rewarded odor, **α**, in both tasks, that is, by making the odor identity and reward association consistent, we isolate the influence of task demands when investigating neuronal responses.

**Figure 1:**
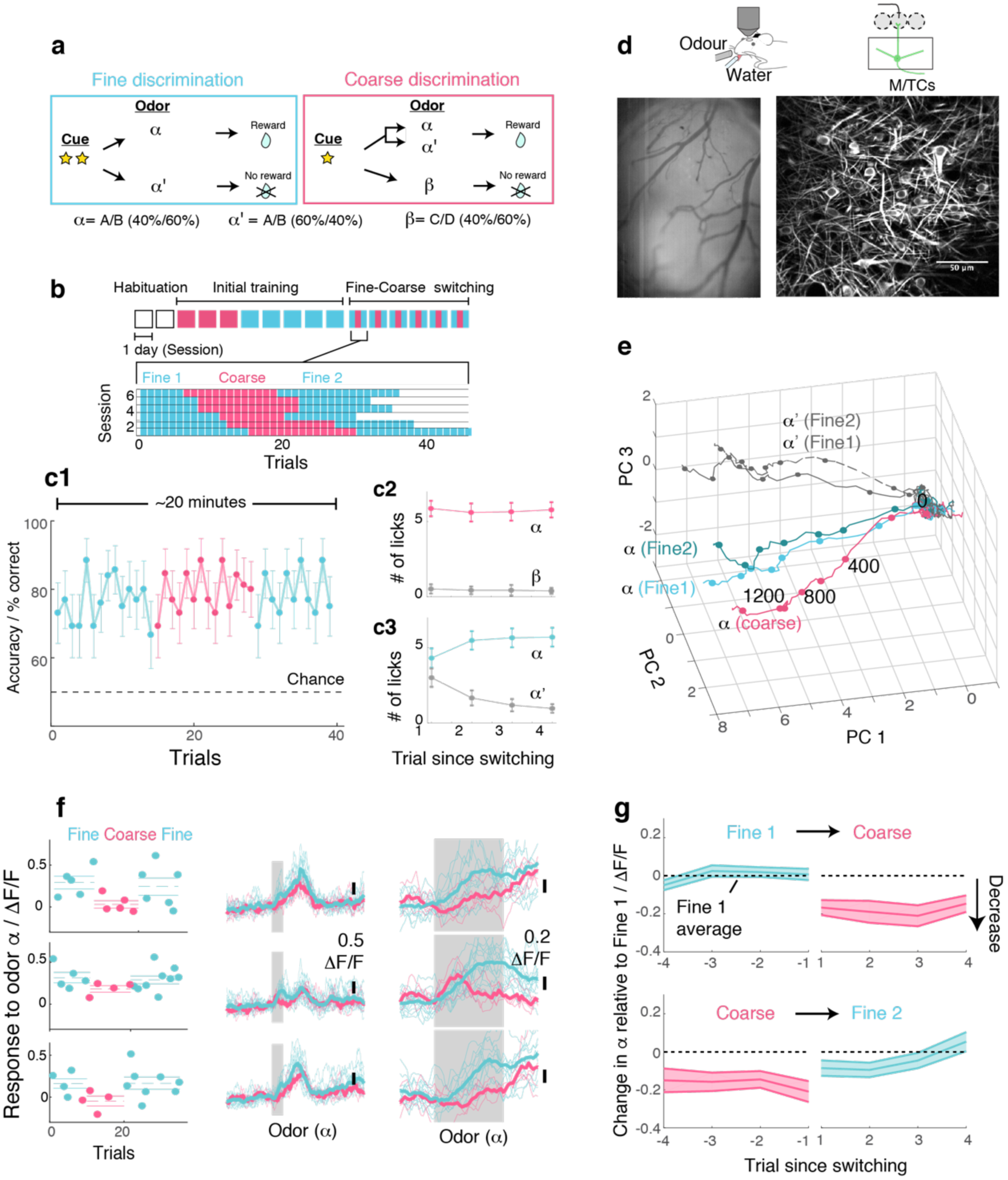
OB output neurons change olfactory representation rapidly and reproducibly with demands. (**a**) Structure of discrimination tasks. (*Left*) Fine discrimination. A trial starts with two flashes of an LED. A rewarded stimulus (α) is a mixture of A and B odors at a concentration ratio of 40:60% (α), associated with a water (reward). In a non-rewarded trial, the A/B mixture is presented at a concentration ratio of 60:40% (α^’^), and no reward is given. (*Right*) Coarse discrimination. A trial starts with one flash of an LED. The rewarded odor is the same A/B 40/60 mixture as in fine discrimination. A non-rewarded stimulus is a mixture of different odors (β). On some rewarded trials, α^’^ odor is presented to assess whether mice generalize across both A/B mixtures. (**b**) Timeline of experiment. Each session lasted ^~^20 minutes, and occurred once a day. *Inset*: Example of switching sessions used for a typical animal (6 sessions shown). Variable epoch length ensures that at least 4 rewarded trials appear per epoch. (**c**) Trial-by-trial average performance across 5 mice for all trials (**c1**), as well as trials around switching (**c2,c3**). Mean and s.e.m. are shown. (**d**) Imaging configuration. GCaMP6f fluorescence from mitral and tufted cells was imaged with a two-photon microscope through a chronic window (*middle*) in head-fixed mice performing task switching. *Right*, an example field of view. Scale bar = 50 μm. (**e**) Trajectories corresponding to the α and α^’^ odors during fine and coarse discrimination, plotted as trajectories in the first three principal components. Pseudo-population response constructed from all ROIs (n = 353, 5 mice). (f) *Left*, Trial-by-trial α odor response (average amplitude during 1 s odor presentation) for example ROIs. Mean and s.e.m. are shown as horizontal lines. *Middle*, Corresponding transients averaged for each task type. *Right*, The same traces as in the middle panel, but zoomed into the odor period. (**g**) Time course of change among significantly modulated ROIs. The response amplitude for an odor relative to response amplitudes during the first fine discrimination epoch (mean ± s.e.m.; n = 42 ROIs, 5 mice).

Following sequential training for coarse and fine discrimination over two weeks, head-fixed mice were trained to switch rapidly between the two tasks within the same imaging session (Fig. 1a,b). A typical session lasted approximately 20-30 minutes, consisting of two fine discrimination epochs (designated “Fine 1” and “Fine 2”) and one coarse discrimination epoch that occurred between the two fine discrimination epochs. This design was used to control for time-dependent effects. In some trials during the coarse discrimination, odor **α^’^** (the 60A/40B mixture) was presented as a rewarded odor (“probe” trials). This modification forced mice to generalize over A/B mixtures (**α** and **α^’^**) during coarse discrimination, and to discriminate between the **α** and **α^’^** odors during fine discrimination. Over the course of four days, on average, mice learn to perform the task switching with high accuracy (Fig. 1c, Supplementary Fig. 2).

To assess the effect of task demands on OB processing, olfactory representation was imaged simultaneously in the principal neurons, mitral and tufted cells (M/TCs), using a two-photon microscope (Fig. 1d). A genetically encoded calcium indicator, GCaMP6f (Chen et al., 2013), was expressed conditionally in M/TCs in the OB by crossing the Ai95D mouse line (Madisen et al., 2015) with the Tbet-ires-Cre mouse line (Haddad et al., 2013). Imaging was accomplished through a previously implanted chronic imaging window (Holtmaat et al., 2009).

To visualize if and how M/TC responses as a whole are modulated by tasks, calcium transients from all regions of interest (ROIs; 353 ROIs from 5 mice) were expressed as pseudopopulation vectors and plotted as trajectories in the first 3 principal components (Fig. 1e). Despite the fact that the odor is identical, the trajectory for the α odor during coarse discrimination lies distinctly away from that for fine discrimination. On the other hand, the trajectories for odor **α** from the first and second epochs of fine discrimination superimpose closely. This indicates that olfactory representation in the OB changes with task, but reversibly.

At the level of individual cells, a subset of MTCs was found to change its responses significantly with task (42/353 ROIs; 17/353 ROIs in shuffle control show significant change). Single-trial analysis revealed that lick and sniff patterns do not explain this task-related change (Supplementary Fig. 3). In fact, when variability arising from sniff patterns was removed through linear regression, a greater proportion of ROIs were found to be significantly modulated by task (56/353 ROIs; Supplementary Fig. 3). Similarly, task-related modulation is not explained by the difference in the stimulus statistics (frequency of A/B mixture presentations), as the modulation is absent in mice anaesthetized with ketamine and xylazine (Supplementary Fig. 4). Importantly, the change is present immediately after task switching (Fig. 1f,g). Among the task modulated ROIs, significant change is observed even in the first trial after mice switch from fine to coarse discrimination (mean change in the α odor response during 1^st^ coarse trial = −0.17 ± 0.04 ΔF/F; p<0.01, t-test for equal means, t-score = −4.3, n = 43 ROIs, 5 mice). Switching back to fine discrimination, responses become comparable to the original amplitudes by the 3^rd^ trial (mean difference in α odor response relative to Fine 1 = −0.05 ± 0.04 ΔF/F; p = 0.26, t-score = −1.2), closely mirroring the recovery time course of behavioural accuracy (Fig. 1c).

Overall, M/TCs tend to increase their responses during fine discrimination (Fig. 2a,b). The increase is particularly pronounced when mice show clear evidence of switching, as assessed by the performance during probe trials (Supplementary Fig. 5).

**Figure 2:**
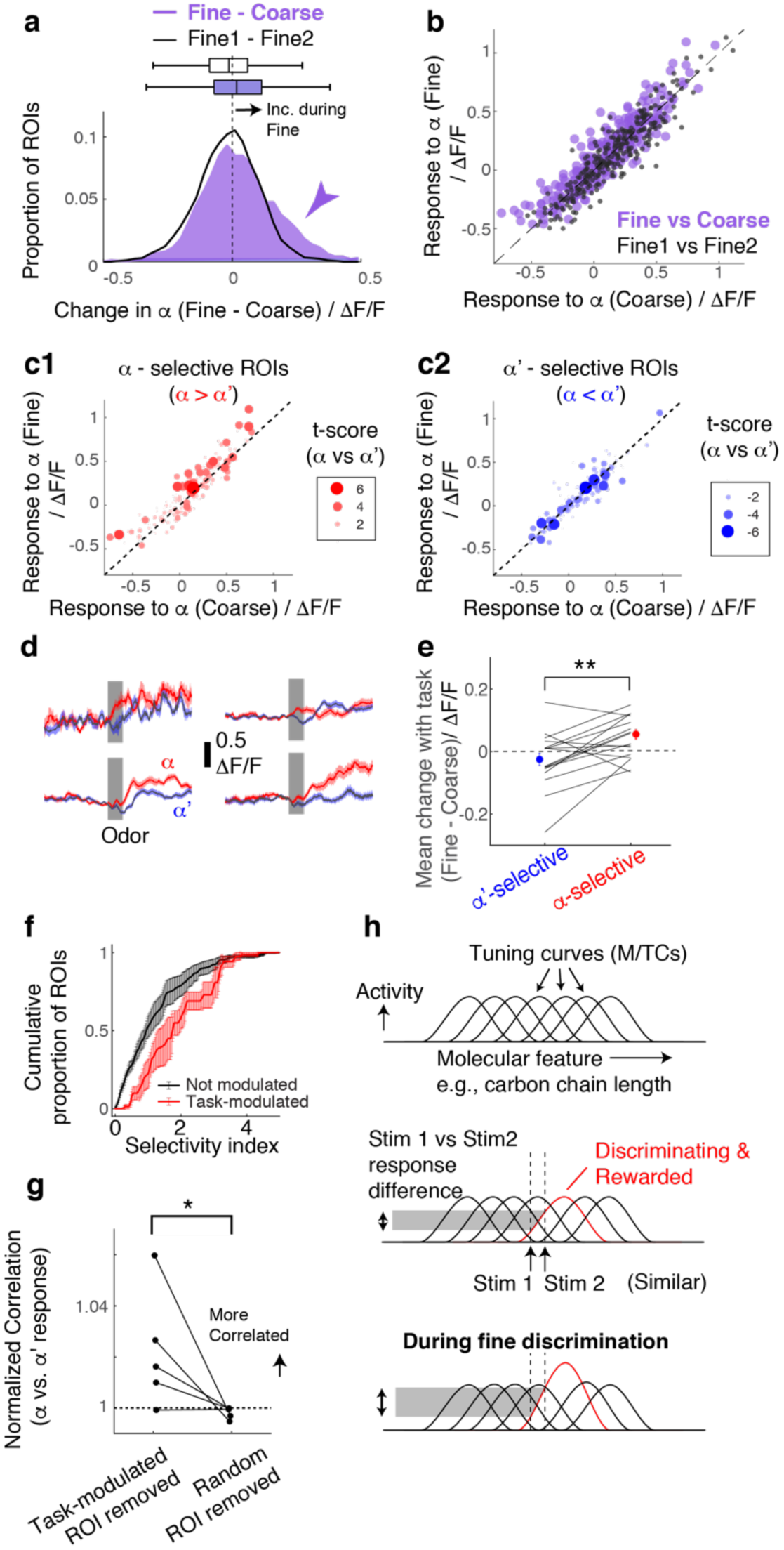
Selective amplification leads to demand-specific enhancement of stimulus decorrelation. (**a**) Histogram of task-dependent change in α responses (response during fine discrimination – response during coarse discrimination; purple). Within-task variability (black line) is shown for comparison. (**b**) Comparison of odor responses during coarse (x-axis) and fine (y-axis) discrimination for all ROIs (purple). Within-task comparison (Fine1 vs Fine2; black points) shown for reference. (**c**) Task-dependent odor response comparison as in (**b**), but separated by odor selectivity. α-selective ROIs (red) are ROIs with t-scores greater than zero, where the strength of the selectivity is indicated by the size of the marker. α^’^-selective ROIs are those with t-scores < 0. (**d**) Average transients in response to odor α (red) and odor α^’^ (blue) from 4 ROIs selective for odor α. Error bar = s.e.m. (e) Summary of task-dependent change for odor α (top) for ROIs grouped by selectivity. Each data point = average for each imaged location. N = 21 sessions, 5 mice. Sessions where probe trials were omitted are excluded from this analysis. (**f**) Distribution of selectivity indices for task-modulated ROIs (red; n = 32 ROIs, 5 mice) and ROIs without task modulation (black; n = 255 ROIs, 5 mice). The selectivity index is the absolute value of the t-score for α-responses vs α^’^-responses imaged during fine discrimination. (**g**) Correlation coefficients between α vs α^’^ odors, where a subset of ROIs is removed and normalized to when a full set of ROIs used. Each coefficient is from ROIs from individual animals, instead of imaging location, due to a small number of ROIs in some imaged locations (n = 5 mice). (**h**) Schematic of finding. MTCs tuned differently for a given “molecular feature,” for example, the carbon chain length, have different tuning curves along this axis. Some MTCs respond differently to two similar stimuli (stim 1 and stim 2), which contribute the most to discrimination. MTCs that are selectively tuned to the rewarded odor are enhanced when mice need to discriminate between the two similar odors, resulting in enhanced discriminability.

According to a previous, longitudinal study, when mice learn to perform fine discrimination, there is an accompanying increase in the fraction of divergent responses among M/TCs (Chu et al., 2016). Accordingly, we assessed whether stimulus selectivity is an important parameter in the dynamic, task-related change observed here. To this end, for each ROI, we measured its selectivity to α vs α^’^ odors using the t-score (or t-statistic), which compares mean response amplitudes (see Methods). Intriguingly, we found that it is the α-selective M/TCs that enhance their responses during fine discrimination (Figure 2c-e; mean change = 0.06 ± 0.02 ΔF/F for alpha selective ROIs; p = 0.008, paired t-test for equal means between α-selective and α^’^-selective ROIs; n = 21 sessions, 5 mice).

As a result, stimulus-selective neurons are over-represented among task-modulated M/TCs (Fig. 2f; p = 0.01, 2-sample K-S test for equal distributions, K-S statistic = 0.33, n = 22 task-modulated ROIs and 201 non-modulated ROIs from sessions with probe trials), suggesting a critical role that modulation plays in discriminating similar stimuli. Consistently, when these, task-modulated ROIs are removed, the population of M/TC responses to α and α^’^ odors became more correlated (Fig. 2g; mean % change in Pearson’s Correlation = 2 ± 1 when modulated ROIs were removed, and 0.2 ± 0.1 when random ROIs were removed; p = 0.04, paired t-test, n = 5 mice). We note that a general increase in response amplitudes alone does not explain our result, as an overall increase in response magnitude in an anaesthetized preparation does not lead to enhanced stimulus selectivity and discriminability (Supplementary Fig. 6).

## Discussion

Overall, we find that rapid, task-dependent modulation of odor responses in the primary olfactory area occurs dynamically to enhance odor representation to suit the behavior. We find that modulation occurs even when the stimulus-reward association is identical, isolating context as the only variable. Concomitantly, discriminability of stimulus representations is altered through selective boosting of responses of discriminating neurons (Fig. 2h).

We also show that changes in the bottom-up input, including sniff patterns and stimulus statistics, are inadequate to explain the task-dependent modulation observed here. How, then, might dynamic and selective modulation arise in the OB? Unusually for a primary sensory area, the OB receives dense feedback projections from olfactory cortices (Boyd et al., 2012; Davis and Macrides, 1981; Markopoulos et al., 2012; Otazu et al., 2015; Rothermel and Wachowiak, 2014) as well as neuromodulatory centres(Macrides et al., 2004; McLean and Shipley, 1987; Steinfeld et al., 2014). Among these, possible pathways for selective amplification include direct excitatory feedback, such as a direct excitatory drive from centrifugal inputs onto M/TCs, as described for the AON (Markopoulos et al., 2012), or modulation of inhibitory neurons via some competitive mechanism(Koulakov and Rinberg, 2011), although we cannot rule out a role for neuromodulators. So far, there is no evidence that olfactory signals carried by the cortical feedback are spatially matched to local (glomerular) olfactory representations (Boyd et al., 2015). Whether cortical feedback to the OB indeed forms a pattern akin to an “attention-field” to enhance relevant signals (Reynolds and Heeger, 2009) will be a key topic of future investigation.

In discriminating similar stimuli, modeling studies suggest that neurons that contribute the most are those that respond differently to the stimuli used. These are most often neurons with a preferred stimulus feature that is shifted slightly away from the stimulus features to be discriminated (Jazayeri and Movshon, 2006). Our results suggest that such neurons may be the target of selective and dynamic modulation to enable enhanced discriminability as needed. The ability to induce different task demands and observe the resultant modulation in a primary sensory area may prove useful to understand how the brain implements solutions that meet ever changing behavioural demands.

## Methods

### Animals/surgery

All animal experiments were approved by the UK Home Office and the institutional veterinarian and ethics committee. All recovery surgery was carried out using standard aseptic technique under isofluorane anaesthesia and carprofen analgesia. Adult male mice (6 – 8 weeks old) from the cross between the Tbet-Cre line(Haddad et al., 2013) and the Rosa26-GCaMP6f line (Madisen et al., 2010) were used. For chronic optical access, a method similar to that of Holmaat et al (Holtmaat et al., 2009) was used. Briefly, a small, round piece of glass (^~^ 1 mm diameter) was cut from a cover slip (borosilicate glass 1.0 thickness) using a diamond knife (Sigma-Aldrich). It was fitted over a craniotomy made above the left olfactory bulb and sealed with cyanoacrylate (Histoacryl, TissueSeal, USA). A custom-made head-plate was attached to the exposed, dried occipital bone using gel superglue and dental cement. Non-steroidal anti-inflammatory treatments (Carprofen, i.p.) were given for 3 days postoperatively. All behavioural sessions began more than 2 weeks after the surgery.

### Olfactometry

Odors were presented using a custom-made flow-dilution olfactometer similar to an earlier design (Fukunaga et al., 2012), except in the control of odor concentrations (Supplementary Fig. 2). Odor concentration was set using precise, pulsatile packets of saturated odor into a 50-mL flask, where it was mixed with a steady flow of background air (1 slpm) in order to eliminate fast transients in odor concentration (time constant =^~^400 ms). The final odor concentration presented to the animal was approximately 0.5-1% of the saturation level. A binary mixture was generated by mixing two streams of odorized air in the same mixing compartment, and concentrations were verified using a photoionization detector for each odor component (where the tested odor was mixed with a blank control that did not give signals; Supplementary Fig. 2). Each odor canister had a large headspace (^~^40 mL) over the odorants to minimise run-down over time. Teflon tubing and air purge (^~^ 5 slpm for 10 seconds) during the inter-trial interval were used to minimize odor contamination.

### Go-No-Go olfactory discrimination

Water access in the home cage was restricted, and head-fixed mice went through habituation sessions before discrimination training commenced. During habituation sessions, animals were head-fixed and were presented water from the port. This stage of training lasted 15-20 minutes a day, until animals were comfortable enough to drink water from the water port. This was typically achieved within two days. Animals were then trained to associate one olfactory stimulus with a water reward (S+ odor) and another olfactory stimulus with no reward (S-odor). The correct response for the S+ trials was to lick the water port for a reward during 2 seconds after the onset of odor presentation, while the correct response for S-trials was to refrain from licking. The reward was a single drop of water approximately 20 uL in volume. The water port served also as a lick sensor (beam break; PM-F25, Panasonic, Japan), which was coated with black silicone externally to prevent light leakage. Respiratory rhythms were monitored continuously through the contralateral naris using a fast mass flow sensor (FBAM200DU, Sensortechnics, Germany). Odors used were ethyl butyrate and eugenol in the rewarded odor mixture, and methyl salicylate and methyl tiglate in the non-rewarded odor mixture used in the coarse discrimination. The sequence of S+ and S-trials was chosen so that no more than three consecutive trials were of the same type, but otherwise random. On each trial, odor was presented for 1 second, and inter-trial interval was 20 seconds. Animals were monitored closely and at no point did animal weight decrease below 80% of the original.

### Imaging

Data in this manuscript were obtained from awake, head-fixed mice engaged in the tasks, as well as anesthetized mice, as indicated, through previously implanted optical windows. Two-photon fluorescence of GCaMP6f was measured with a custom-fitted microscope (INSS, UK for M/TC imaging) and high-power laser (930 nm; Insight DeepSee, MaiTai HP, Spectra-Physics, USA) at depths 50-400 μm below the surface of the olfactory bulb. Images from a single plane were obtained at ^~^30 Hz with a resonant scanner. Each day, the stage was zeroed at a reference location, guided by the surface blood vessel pattern using epifluorescence, and sampled from an area not overlapping with previous sessions (based on the co-ordinates and field view, and confirmed post-hoc by eye).

### Analysis

Images were analysed using custom-written routines in Spike2 (Cambridge Electronic Design, UK), Matlab (Mathwarks, USA) and macros in Fiji (ImageJ), run on a PC (32GB RAM, 10 x 2.2 GHz Intel Xeon processors).

<u>*ROI detection*</u>. From an average of ^~^1000 frames for each imaging plane, somata were manually delineated using an ROI manager (ImageJ). Selected oval regions were confined within the perimeter of the soma, and somata with filled nuclei were excluded. Based on the method by Kerlin et al(Kerlin et al., 2010) (Supplementary Fig. 6), the contribution from out-of-focus fluorescence was approximately 30%. <u>*Odor response amplitude.*</u> Unless otherwise stated, the average odor response was obtained from ΔF/F values during the first 1 s of odor presentation (after the onset of valve opening). Results are qualitatively similar when measurements are shorter time windows, but noisier, especially for single-trial analyses. <u>*Similarity of S+ vs S-representations.*</u> For each pair of responses per imaging location, the relative fluorescence change (ΔF/F values) for all ROIs in the field of view was treated as vectors and Pearson correlation coefficient was calculated. <u>*Odor response.*</u> Significant changes were those in which the magnitude of odor-evoked change exceeded 3x the standard deviation of baseline activity. <u>*Task-dependent change.*</u> To determine if a task-dependent change was statistically significant, evoked response amplitudes for S+ trials for fine and coarse discriminations were tested for the same distributions (two-sample t-test for no difference, two tailed). <u>*Principal component analysis.*</u> The method is based on Niessing and Friedrich(Niessing and Friedrich, 2010). Briefly, for each ROI, GCaMP6f transients (ΔF/F values) from each condition (odor and task) were concatenated and principal components obtained using the Matlab function *pca*. Original data were projected on the new components (first three principal components) to obtain trajectories. <u>Stimulus selectivity (T-score)</u>. For each ROI, odor response amplitudes were obtained from α trials and α^’^ trials. These were compared using the Matlab function for the two means, namely, the two-sampled t-test (ttest2) to obtain a t-score (t-statistic). The T-score was used instead of the commonly used z-score because of the small number of trials. <u>Selectivity index</u> was defined as the magnitude (absolute value) of the t-score. <u>ROI removal and resultant change in stimulus correlation</u>. For each animal, average odor response from ROIs (all imaging sessions) were expressed as one vector. From this, task-modulated ROIs were removed and Pearson’s correlation coefficient for α- and α^’^-responses was calculated. This value was compared against the correlation coefficient obtained from the full set i.e., change in correlation = coefficient (ROI removed) – coefficient (full set))/coefficient (full set). For removing random sets of ROIs, a random subset, comprising the same number of ROIs as the task-modulated set, was selected using the randperm function in Matlab. Then the change in correlation was calculated as above. <u>*Linear decoder analysis.*</u> Matlab function fitcdiscr was used to fit a linear classifier. For each dataset (each imaging session), training data comprised the population of M/TC responses from 2/3 of the trials during the fine discrimination block. The other 1/3 of the trials was used as a test set to predict the label (S+ vs S-odor). % accuracy is the % of trials from the test set that matched the correct label. <u>*Behavioral performance.*</u> Animal lick responses were analysed. For S+ and S-trials, correct responses were the presence and absence of licking during a window 1-2.5 seconds after odor onset, respectively. Incorrect responses were the absence and presence of lick responses for S+ and S-trials in the same window, respectively. The number of correct trials was expressed as the % of all trials.

## Acknowledgement

The authors thank Martyn Stopps for help with the olfactometer design, Charly Rousseau and Nicholas Burczyk for technical advice, Molly Strom and Troy Margrie for reagents, and Kevin Franks, Alexander Fleischmann, Rebecca Jordan, and Steven Aird for helpful comments on the manuscript. This work was supported by Okinawa Institute of Science and Technology Graduate University, the Francis Crick Institute which receives its core funding from Cancer Research UK (FC001153), the UK Medical Research Council (FC001153), and the Wellcome Trust (FC001153); by the UK Medical Research Council (grant reference MC_UP_1202/5); and by the DFG (SPP 1392). AS is a Wellcome Trust Investigator (110174/Z/15/Z).

## Author contributions

IF and AS conceived the project. IF designed the experiments. AK and IF performed the experiments and analysed the data, with input from AS. IF and AS wrote the manuscript with input from AK.

## Supplementary Figures

**Supplementary Figure 1:**
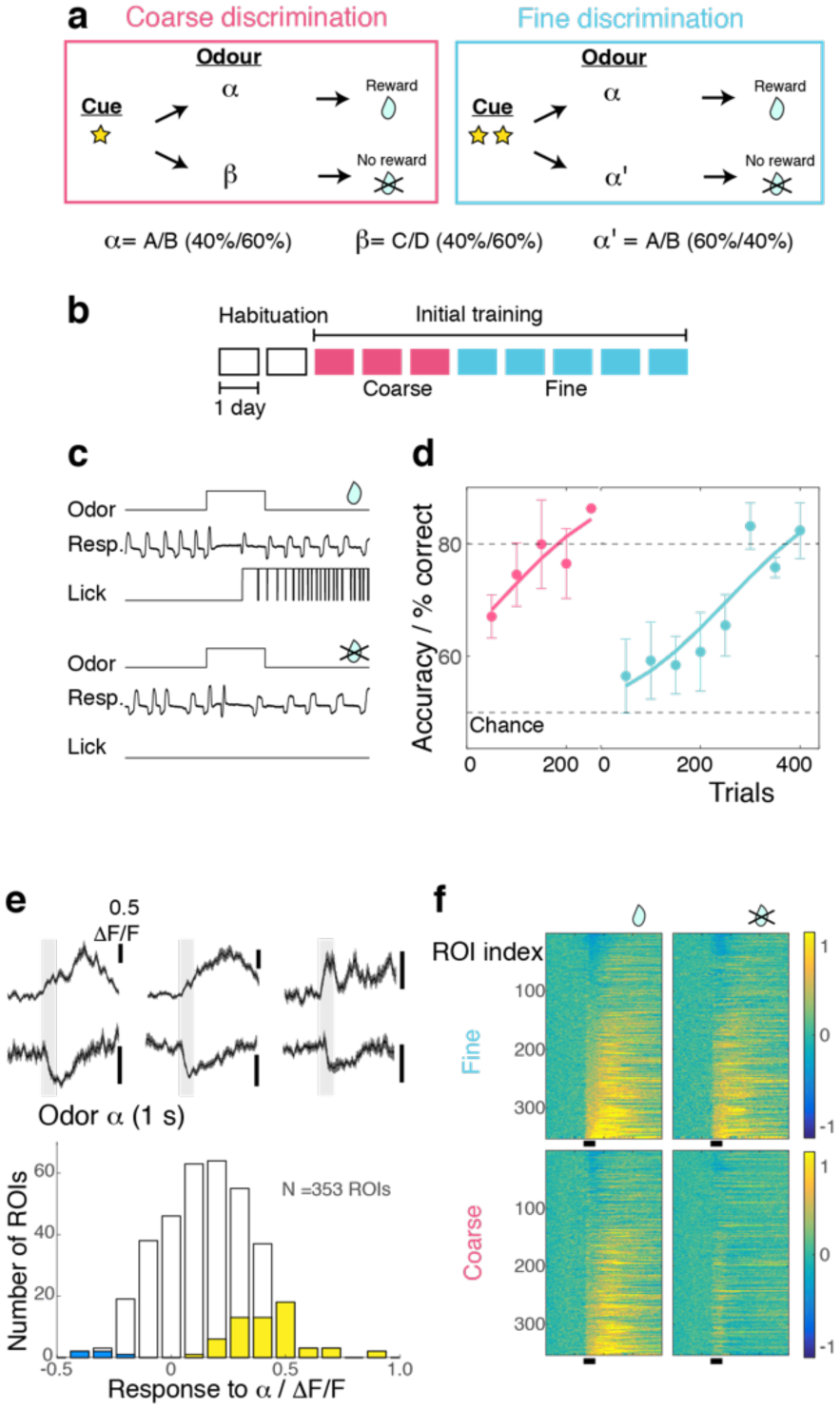
Method for imposing different behavioural task demands. (A) Structure of discrimination tasks. *Left*, Fine discrimination. A trial starts with two flashes of an LED. The rewarded stimulus (α) is a mixture of A and B odors at a concentration ratio of 40:60% (α), associated with a water reward. In a non-rewarded trial, the A/B mixture at a concentration ratio of 60:40% (α’) is presented, but no reward is given. *Right*, Coarse discrimination. A trial starts with one flash of an LED. The rewarded odor is the same as in fine discrimination. A non-rewarded stimulus is a mixture of different odors (β). In some rewarded trials, α’ odor is presented to assess whether mice generalize across both A/B mixtures. (**C**) Examples of correct behavioural responses for rewarded trials (top; correct hit) and for non-rewarded trials (bottom; correct rejection). An odor is presented for 1 s. and a reward is delivered 3 s after onset of the odor. (D) Learning rates for head-fixed mice used in the experiment (n = 5 mice). (**F**) *Above*. Examples of excitatory and inhibitory responses evoked by S+ odor (gray bar, 1 s). *Below*. The distribution of responses evoked by S+ odor. Yellow and blue bars = significantly excitatory and inhibitory responses, respectively. (**G**) Calcium transients as color maps in response to rewarded (*left*) and non-rewarded (*right*) odors for the tasks indicated. ROIs are sorted by the S+ response amplitude. n = 353 ROIs, 5 mice.

**Supplementary Figure 2:**
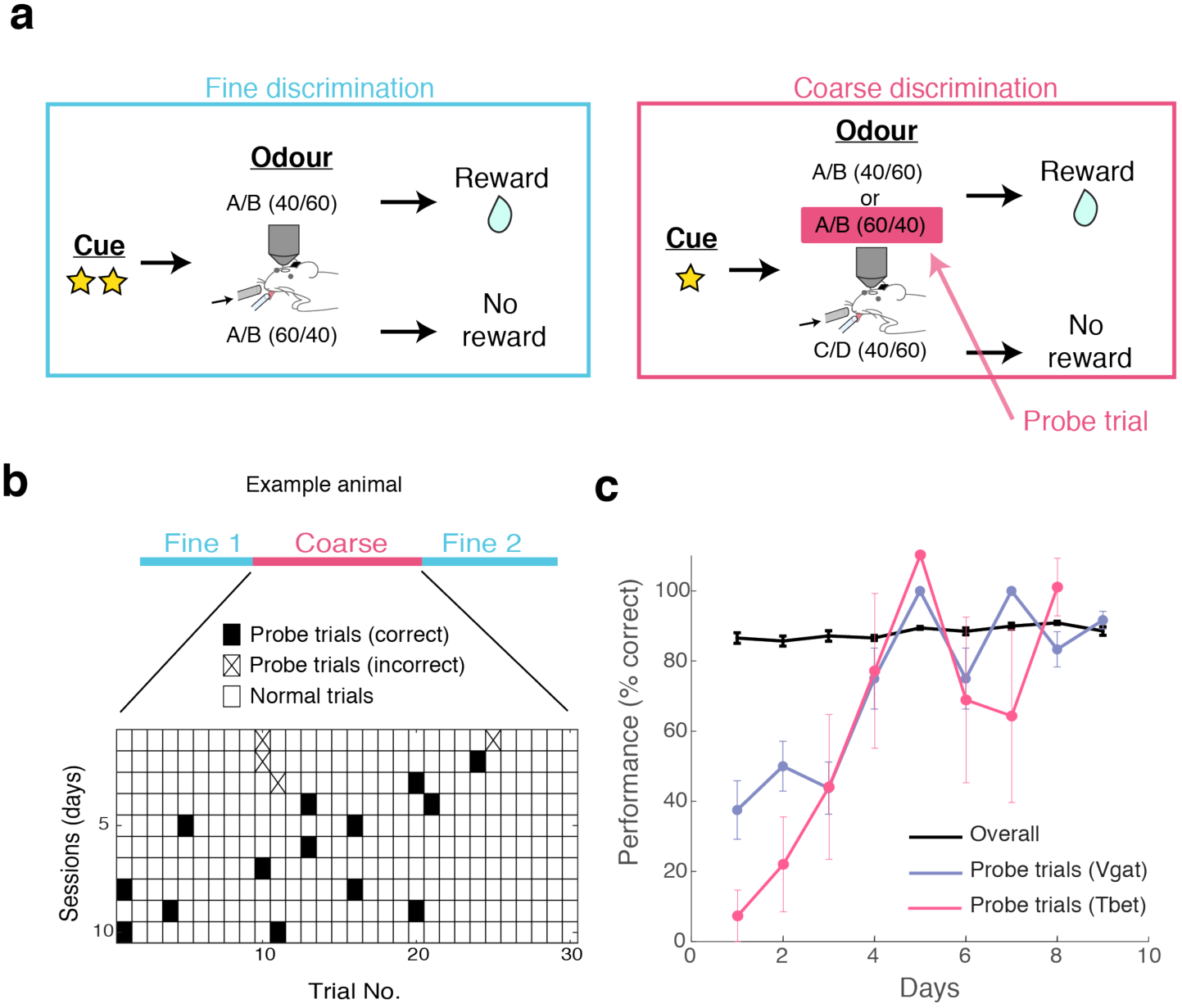
Task-switch learning and probe trials. (A) Behavioural paradigm for task switching. Mice are trained to perform fine discrimination blocks before and after a coarse discrimination block within one session. The design of the coarse discrimination is such that mice are forced to generalise closely related odours. On a given rewarded (S+) trial, the odour was either A/B in a 40/60 mixture or A/B in a 60/40 mixture. This latter stimulus appears as the non-rewarded odour in the fine discrimination session. Therefore, the A/B (60/40) trial during the coarse discrimination block serves as a test to see whether mice genuinely perform tasks differently, so they are termed “Probe trials.” (B) A typical example showing the performance of a mouse during the probe trials over several days (1 session per day). Trials during coarse discrimination are shown as tiles. Probe trials with an incorrect response (i.e., no lick) are marked with a cross, while those with a correct response are shown as filled tiles. (C) On average, mice learned to perform task-switching in 4 days, as seen in the performance accuracy in these probe and overall trials. N = 4 mice for Vgat-ires-Cre experiment and 5 mice for Tbet-ires-Cre x Ai95D experiment (average of 1.6 ± 0.4 trials per mouse per session).

**Supplementary Figure 3:**
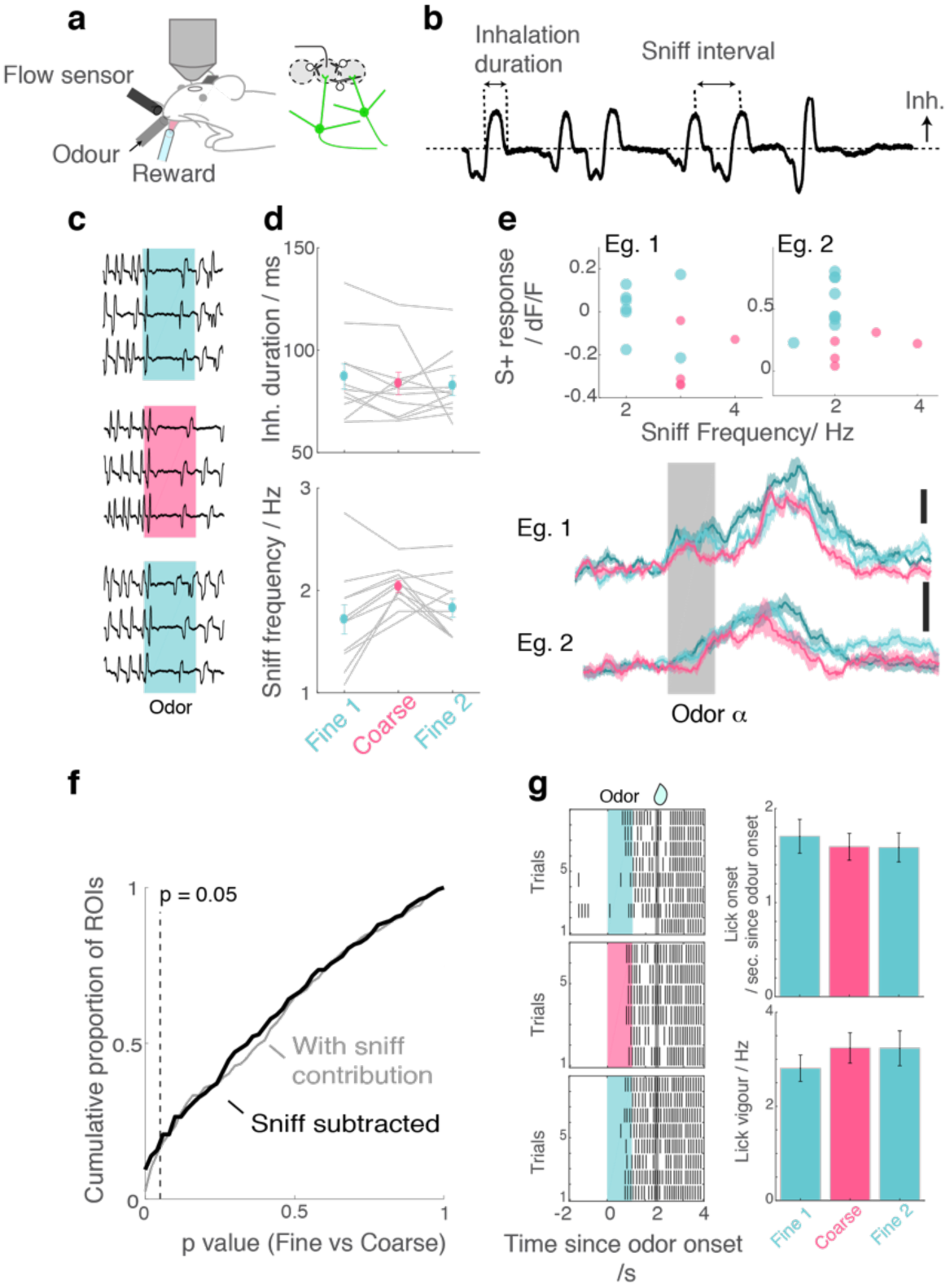
Movement and sniff patterns do not contribute to task-dependent differences. (**A**) Experimental setup. A flow sensor was placed in close proximity to the right nostril to monitor respiratory rhythms while the odor was presented to the left nostril. A lick sensor was a small beam break sensor placed around a water delivery port. (B) Explanation of sniff parameters measured. (C) Example of respiratory traces around the time of odor presentation. Inhalation is plotted upwards. (**D**) Summary of sniffing patterns showing the average duration of inhalation (*above*) and the number of inhalations taken during the 1 s odor presentation (*below*). (**E**) Examples of dependence on sniff: odor-evoked response amplitudes of two example ROIs against sniff frequency for individual trials during fine discrimination (turquoise) and coarse discrimination (magenta). Average transients for the same ROIs (right). (**F**) Distribution of p-values where dependency on sniff is eliminated thorough linear regression (fitted for rewarded odor responses during fine discrimination) in black, and overall p-value distribution where sniff contribution is not subtracted (gray). (**G**) Lick patterns following rewarded odor presentation. *Left*, Examples from one imaging session. Ticks represent occurrence of licks. Each tick represents the onset of licking, relative to the onset of odor presentation. *Right*, Summary showing the timing of the first lick after odor onset (*above*) and frequency of licking for a 3-second window before reward delivery (*below*).

**Supplementary Figure 4:**
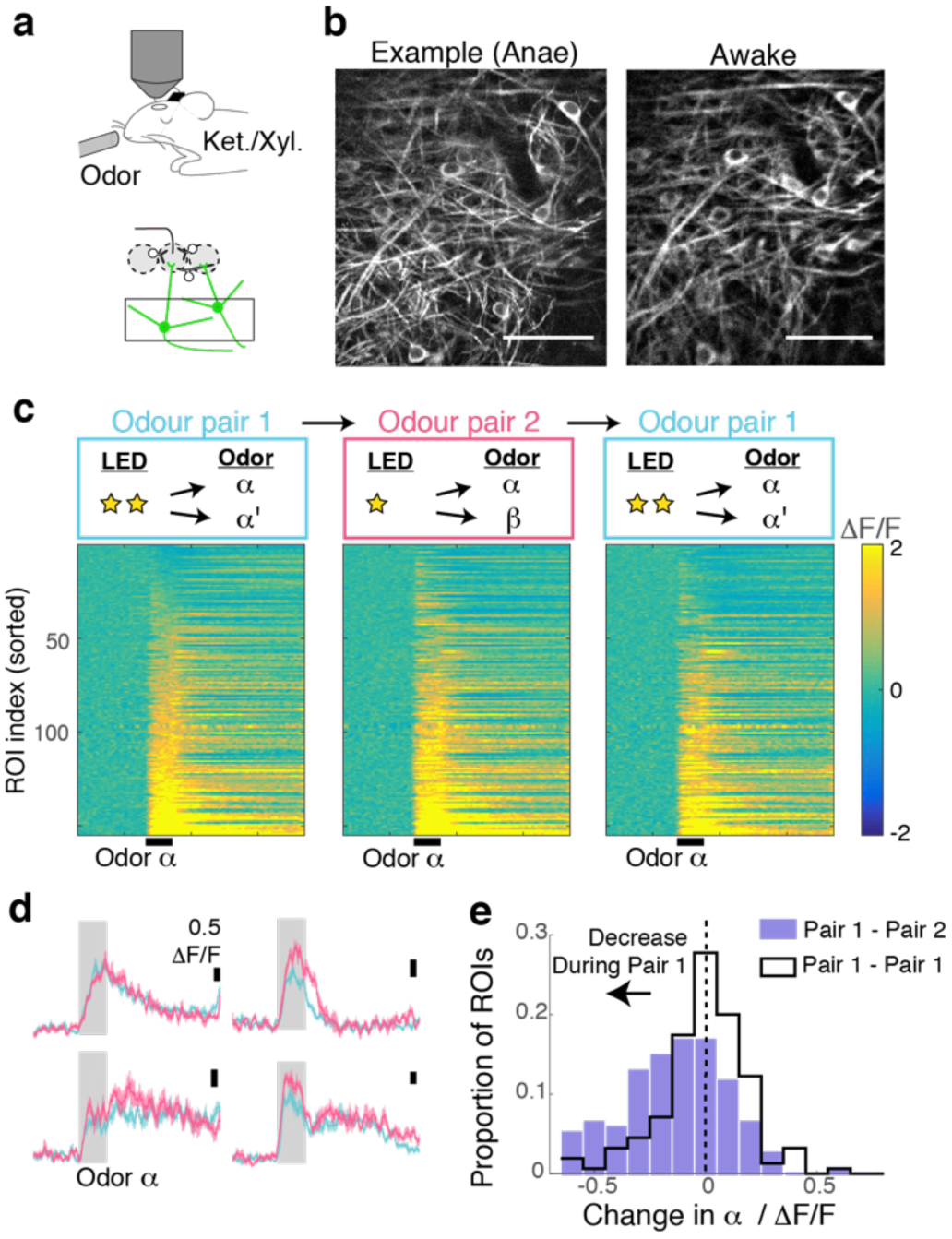
Task-dependent modulation observed during behavior is absent under anaesthesia. (**A**) After the behavioural experiments, mice were anaesthetized with ketamine and xylazine (i.p.). (B) Example fields of view from an example animal show the same location imaged during anaesthesia (left) and during behavioural experiment (right). Scale bar = 50 μm. (C) Calcium transients for 155 ROIs imaged, in response to the same rewarded odor. (D) Average transients from 4 example ROIs in response to the rewarded odor during fine (turquoise) and coarse (magenta) stimulus presentations. Scale bar = 0.5 ΔF/F. (E) Histogram of task-dependent change in α odor responses (response during fine discrimination – response during coarse discrimination).

**Supplementary Figure 5:**
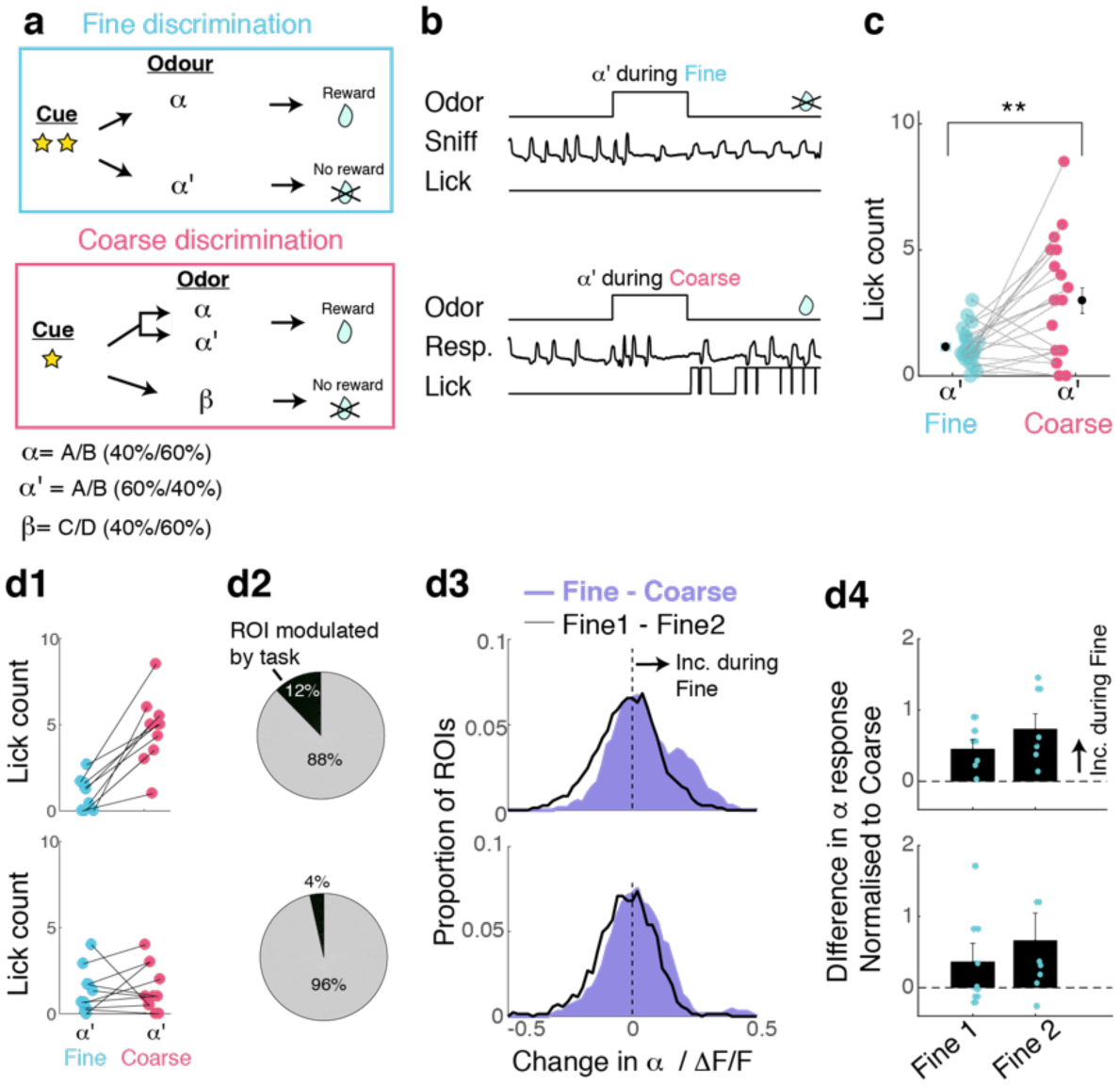
Task-related change is prevalent in sessions with clear evidence of switching. (A) Task structures as before. (B) Example of trials where a 60A/40B mixture (α’ odor) was presented as a non-rewarded odor during fine discrimination (above) and as a rewarded odor during coarse discrimination (below). (C) Summary comparison of lick response for the same odor but during different tasks. (D) Analysis of data from sessions in which mice switched vs those in which they did not. (D1) Definition of switching based on licking pattern following α’ odor presentation. Above: sessions in which mice switched, defined by a significant increase in licking during α’ trials during coarse discrimination. N = 10 imaging sessions, 5 mice. Below, Definition of sessions in which mice did not switch, where licking patterns during α’ odor trials do not differ during fine and coarse discrimination. N = 11 imaging sessions, 5 mice. (D2) Proportion of ROIs that are modulated by task for sessions with evidence of behavioural switching (above) and sessions with no such evidence (below). N = 146 ROIs and 141 ROIs, respectively. (D3) Histogram of change in α odor response amplitudes, across task (Fine - Coarse discrimination; purple) and within task (Fine1 – Fine2, black line). N = same as in D2. (D4) Task-dependent change in a response amplitudes summary representing individual imaging sessions (N = 10 for sessions with behavioural switching, n = 11 for sessions with no behavioural switching).

**Supplementary Figure 6:**
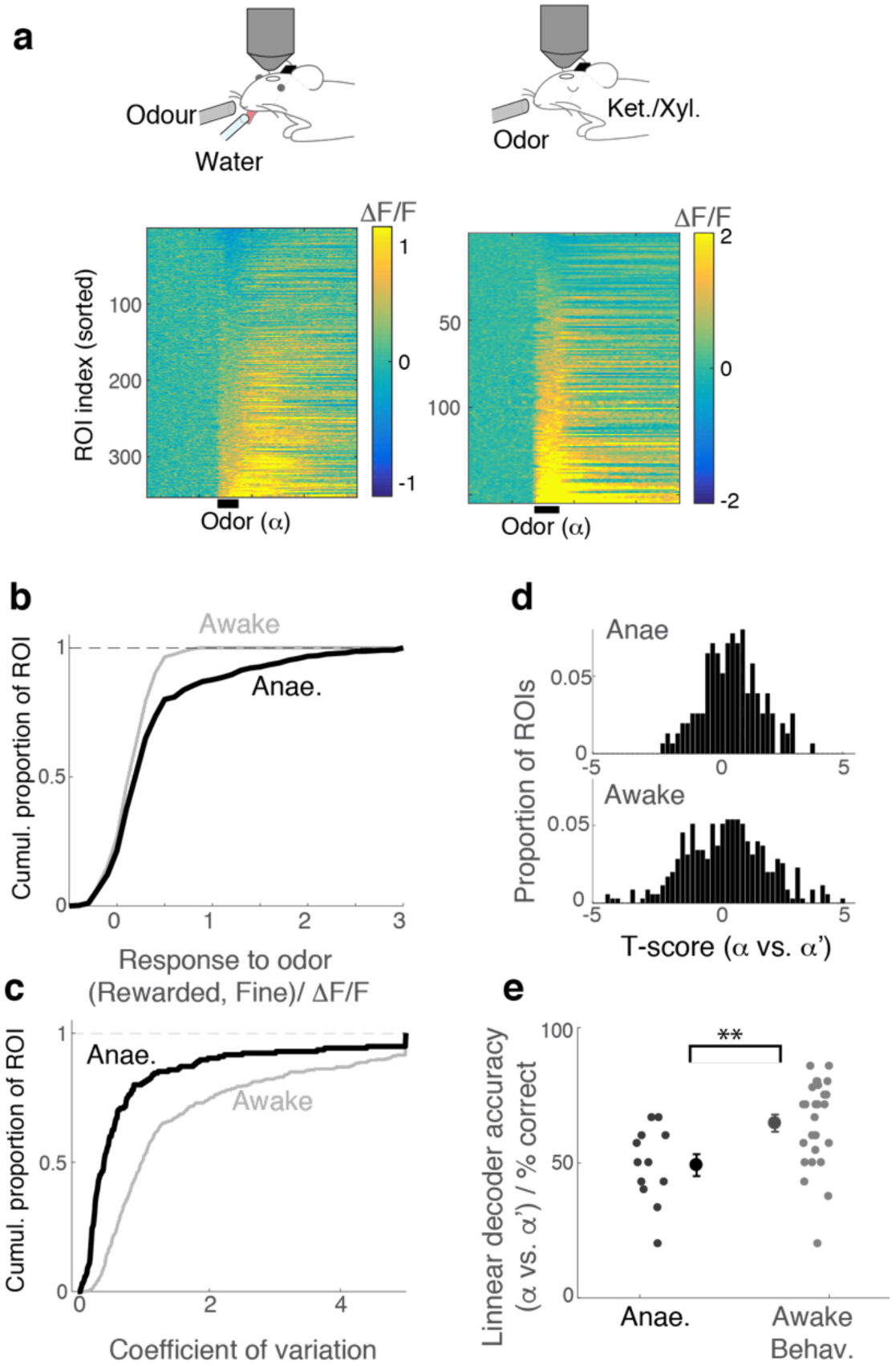
Responses are more reliably discriminated in behaving mice. (A) GCaMP6f fluorescence change is represented in a color map for 353 ROIs imaged from awake, behaving mice (26 imaging sessions, 4 mice). Responses are from mice performing the fine discrimination task. (B) Following behavioural experiments, mice were anaesthetized with ketamine/zylazine and M/TCs imaged (12 imaging sessions, 3 mice; from Fig. 6). Responses are from block 1, where odor α and odor α’ were presented closely in time. (C) Comparison of cumulative histogram of response amplitudes for odor α (average during 1 second after valve opening). (F) Comparison of t-score distribution (α odor response vs α’ odor response amplitudes) for ROIs imaged from anaesthetized (above) and awake, behaving (below) mice. (g) For each imaged location, a linear classifier was trained on a subset of α- and α’ - trials. Its performance for test trials is shown as % correct. (average accuracy = 49.1 ± 4 % correct for anaesthetised vs. 64.6 ± 3.0 % correct for awake, behaving case; p = 0.006, Wilcoxon rank sum test; n = 12 imaged locations, 3 mice for anaesthetized mice vs. 26 locations, 4 mice for awake).

**Supplementary Figure 7:**
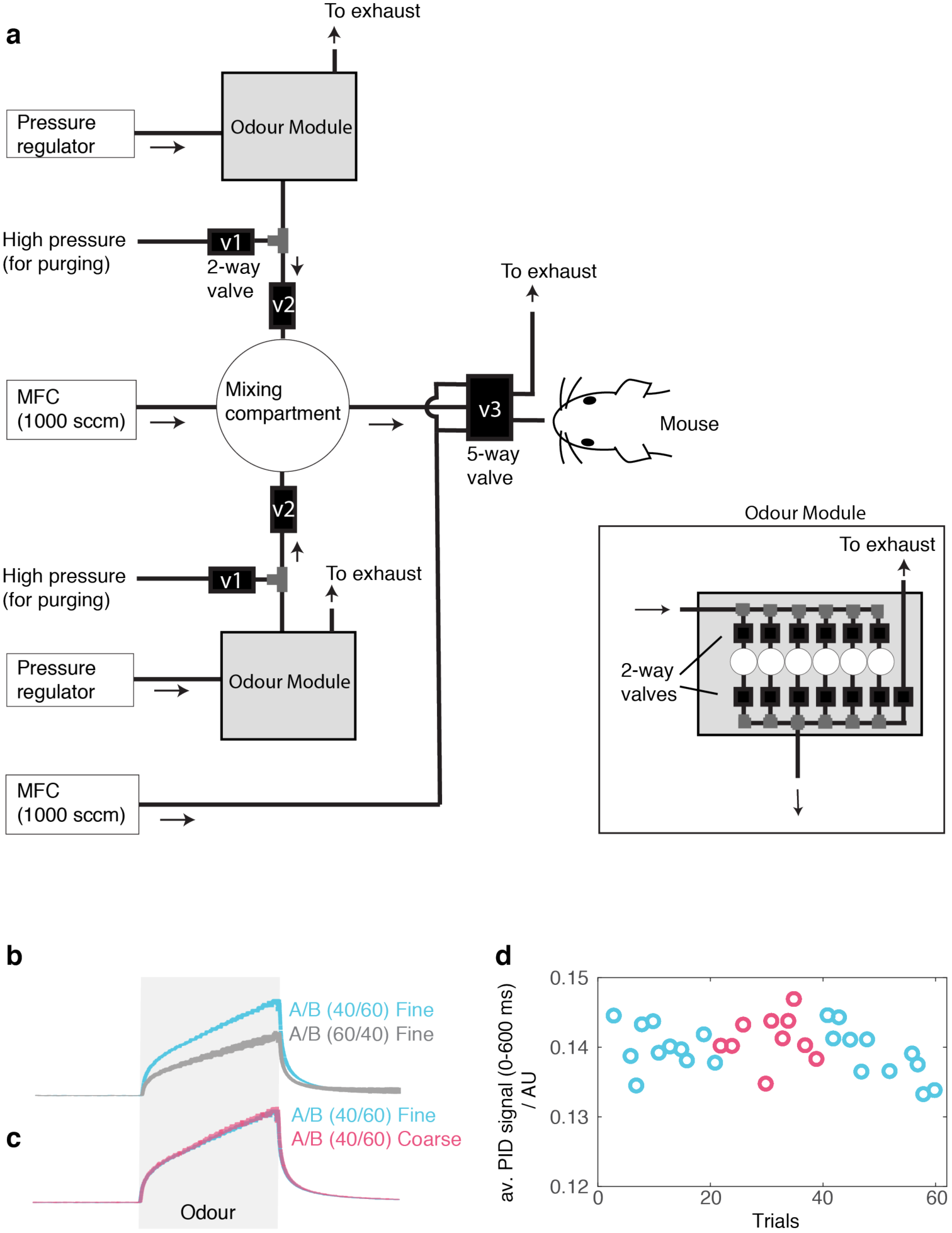
PID signals show stable odour pulses. (A) Olfactometer design for mixing two odorant streams at desired concentrations for generating binary mixtures. For each odorant, filtered and regulated air is passed through a selected odour canister by opening a pair of two-way solenoid valves (Odour module). Odorized air is passed through a second, fast solenoid valve (“v2”) in a pulsatile manner. Pulses of odorized air are mixed in an enlarged compartment (a 50 ml flask) to remove transients. The odour concentration can be set by controlling background air flow and odorized pulse parameters (duration and frequency of rectangular pulses). Mixed air passes through thin PTFE tubing (O.D., 3.2 mm) and is presented to the animal when the 5-way solenoid valve is actuated. (B) For characterizing the stimulus profile, a photo-ionization detector was placed in front of the odour port, in place of the mouse. Instead of the usual binary mixtures, the test used a mixture of 1 odorised stream and 1 blank stream (passing air through an empty odour vial) mixed in the same way to test the concentration of each component separately. PID signals from trials where the composition of the tested odour is 60% (cyan) vs 40% (gray), corresponding to the composition of an odour “A” in S+ and S-trials, respectively. (C) Odour concentrations (PID signal) measured during coarse and fine discrimination for the switching paradigm are overlaid. Mean +-standard error of the mean shown (^~^15 trials each). (D) Time-course of odour concentrations to test the depletion level during a task-switch session. The inter-trial interval was approximately 20 seconds. Data shown are the average PID signal during the first 600 ms of odour period.

**Supplementary Figure 8:**
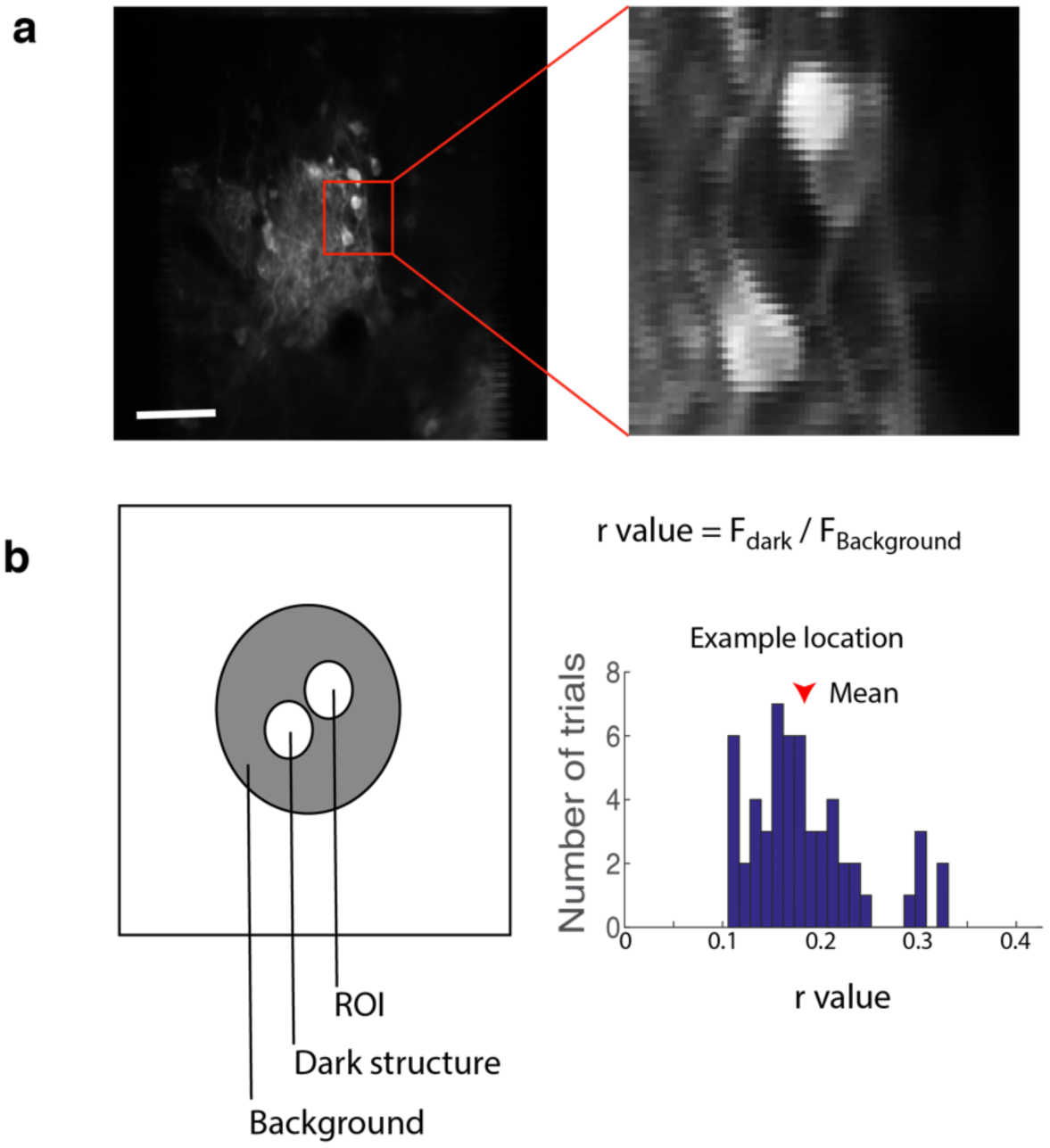
Estimation of contribution from out-of-focus fluorescence. The method is based on Kerlin et al. Briefly, to estimate how much out-of-focus fluorescence contributes to fluorescence within an ROI, the fluorescence of a neighbouring dark object (F_dark_; blood vessel or unstained structure) and the fluorescence of the area surrounding the two objects (ROI and the dark object) are compared. The ratio F_dark_/F_background_ is the estimated fractional contribution from out-of-focus fluorescence.

